# Large shifts among eukaryotes, bacteria, and archaea define the vertical organization of a lake sediment

**DOI:** 10.1101/057117

**Authors:** Christian Wurzbacher, Andrea Fuchs, Katrin Attermeyer, Katharina Frindte, Hans-Peter Grossart, Michael Hupfer, Peter Casper, Michael T. Monaghan

## Abstract

Sediments are depositional areas where particles sink from water columns, but they are also microbial hotspots that play an important role in biogeochemical cycles. Unfortunately, the importance of both processes in structuring microbial community composition has not been assessed. We surveyed all organismic signals of the last ca. 170 years of sediment by metabarcoding, identifying global trends for eukaryotes, bacteria, archaea, and monitored 40 sediment parameters. We linked the microbial community structure to ongoing and historical environmental parameters and defined three distinct sediment horizons. This not only expands our knowledge of freshwater sediments, but also has profound implications for understanding the microbial community structure and function of sediment communities in relation to future, present, and past environmental changes.

## Introduction

The continuous deposition of organic and inorganic particles to sediments is an important process in all aquatic ecosystems. Approximately one-third of the terrestrial organic matter (OM) that enters freshwater is sequestered in sediments [1], although the total amount of OM that reaches the sediments is much greater than the amount that is actually sequestered [2]. This is because microbial activity is responsible for the cycling of carbon, including methane emission [3]. In lake sediments, newly settled OM is rapidly recycled and subsequently transformed into secondary compounds in a distinct uppermost sediment zone of high heterotrophic activity [4,5]. This is thought to lead to the structuring of microbial communities along steeper environmental gradients and narrower vertical sequences of electron acceptors in freshwater as compared to marine sediments [6], presumably due to the higher OM content at the sediment surface. The nature of this gradient influences the carbon, nitrogen, and sulfur cycles [7–9] and potentially affects the microbial community structure [10].

In contrast to the wealth of studies on marine sediments (cf. the 65 studies of [11]), few studies have examined the vertical microbial community structure of freshwater sediments (e.g., [12–16]). This community of sediment microbes was thought to be dominated by bacteria, together with a smaller fraction of methanogenic archaea (reviews in [6,17,18]). However, this view has been challenged by the recent discovery of an abundance of non-methanogenic archaea in marine sediments [19]. They are assumed to be adapted to low-energy environments, and at least one lineage seems to be specialized in inter alia amino acid turnover [19]. This discovery has led to a greatly revised perception of microbial communities in marine sediments, where archaea appear to be as abundant as bacteria and increase in relative abundance with sediment depth [11]. Data on sediment archaea in freshwater are scarce, and the causes of the significant variation observed among studies remain largely unknown (e.g., [14,20–22]).

Overall prokaryotic activity, biomass, and cell numbers decrease with depth in many freshwater and marine sediments, even in the uppermost layer (e.g., [23,24]). Nonetheless, several studies report similar vertical proportions of active cells and find no accumulation of dead cells in deeper sediments [5,25]. Despite the continuous presence of vegetative cells and resting stages, recent studies of marine systems indicate that the majority of microbial cells in energy-deprived horizons consist of microbial necromass [26,27] and the proportion of living organisms decreases with the increasing age of the sediment. The vertical, progressive OM transformation and the depletion of electron acceptors may eventually lead to an extreme low-energy environment in deeper sediment layers and very slow cell turnover rates, as suggested for sub-seafloor sediments [28]. One of the basic mechanisms thought to explain the vertical distribution of microbes is a one-way input of new microbial communities attached to sinking OM. This leaves us with two simplified, competing structural models: (i) The sediment community is exclusively colonized from the surface, thus the community originating from the water column has a fully nested structure that gradually turns from a complex and rich community at the surface to a structure dominated by progressive cell death with increasing sediment depth. (ii) The sediment community at different depth layers is structured by niche specialists that are well adapted to the specific environmental (redoxÒ, OMÒ, metalÒ, nutrientÒ, etc.) conditions (see [29] for a theoretical framework).

The decomposition rate of settled or buried pelagic dead and living organisms can be assumed to depend primarily on the activity of the indigenous microbial community rather than on other chemical processes (e.g., depurination; [30]). Thus, the decrease in sediment DNA reported for freshwater sediment profiles (e.g., [31]) is likely to be a result of the degradation of nucleic acids from dead organisms, particularly eukaryotes, whose biomass strongly decreases with depth. As a result, the decomposition of buried organisms should be a function of the active community, and yet these organisms are in a constant transition of becoming buried themselves over time. *Vice versa*, varying rates of sedimentation over decades will influence the active microbial community by, for example, shifting the redox gradient. It is known these changing (historical) lake conditions are recorded in lake sediments both as DNA signals and as chemical sediment parameters (e.g., [31]).

An important question that remains is how the decomposition processes within the sediment redox gradient are related to the burial of OM, eukaryotes, and prokaryotes [32,33]. Here, we provide a complete biogeochemical and microbial community analysis of a sediment profile of the meso-oligotrophic hardwater Lake Stechlin in northeast Germany. Our aims were to evaluate (1) the extent to which sediment habitats maintain a one-way hierarchical structure, as suggested by the one-way input of organic matter, (2) how “present” and “past” sediment parameters influence the overall community structure (Box 1), and (3) whether recent novel findings regarding the importance and vertical patterns of marine archaea [11] can be transferred to freshwater sediments. We took four replicate 30 cm deep sediment cores from ca. 30 m water depth. ^137^Cs dating indicated the cores include sediments deposited over the past 150 to 170 years. We examined the chemical conditions and microbial community structure of the cores and simultaneously assessed the vertical community composition of eukaryotes, bacteria, and archaea.

#### Box 1. *Definition of “present” and “past” sediment parameters*

We define *present* parameters as the principal components of all context data derived from a) pore water analysis, which indicates that chemical gradients are caused by the consumption and production of ongoing biological processes (e.g., sulfate and methane), as well as from b) directly measured parameters of microbial activities (e.g., bacterial protein production). The present parameters are therefore an expression of recent microbial processes.

*Past* parameters are the principal components of conservative parameters, which once introduced into the sediments will not change significantly and are therefore an expression of the lake’s history (e.g., heavy metals). Here, we also categorize the total amount of elemental carbon, nitrogen, hydrogen, and sulfur as mainly conservative parameters. The past parameters are therefore an expression of historical changes.

## Results

The sediment cores were black in color, with no visible lamination, and they had a high water content (93—97%). Macrozoobenthos was visually absent, and metabarcoding (see below) detected only the presence of nematodes (Figure S1). A consistent pattern across all four sites was the exponential increase of dissolved refractory carbon with sediment depth (Fluorescence Index [*FI*] = 1.69—2.01; Figure 1), indicating the enrichment of fulvic acids [34].

Mean prokaryotic cell numbers were 1.8 ± 0.5 · 10^9^ per ml of wet sediment and were higher in the uppermost sediment horizons. Bacterial biomass production (BPP) (range: 0—282 *μg · ml^−1^d^−1^*) decreased rapidly with depth, approaching zero below 10 cm. Total DNA concentration (range< 0.3—17.6 *μg · ml^-1^* sediment) decreased exponentially with depth and was negatively correlated with FI(*r* =—0.886). DNA half-life was inferred to be *t*_1/2_ = 22 *a* (corresponding to 5.4 *cm*; f (*DNA*) = 13.9 · *e*^−0′128*x*^, *r*^2^ = 0.81). The RNA content of the sediment was lower than DNA at all depths, with DNA:RNA ratios ranging from 2.3 - 20.8 (Figure 1). The sediment exhibited a typical electron acceptor sequence with a mean oxygen penetration depth of 4.6 *mm* (SD 1.4; Figure 1). Nitrate and nitrite were depleted rapidly, sulfate approached a constant minimum concentration after 5 *cm,* soluble reactive phosphorous (SRP) and 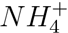 increased with sediment depth, *N*_2_O gas was not detected, *CH*_4_ increased linearly with depth, and *CO*_2_ exhibited minima at the surface and at a depth of 10 cm (Figure 1). Detailed vertical resolution profiles of electron acceptors and profiles of all measured parameters can be found in the supplemental material(Figure S2).

**Figure 1.**
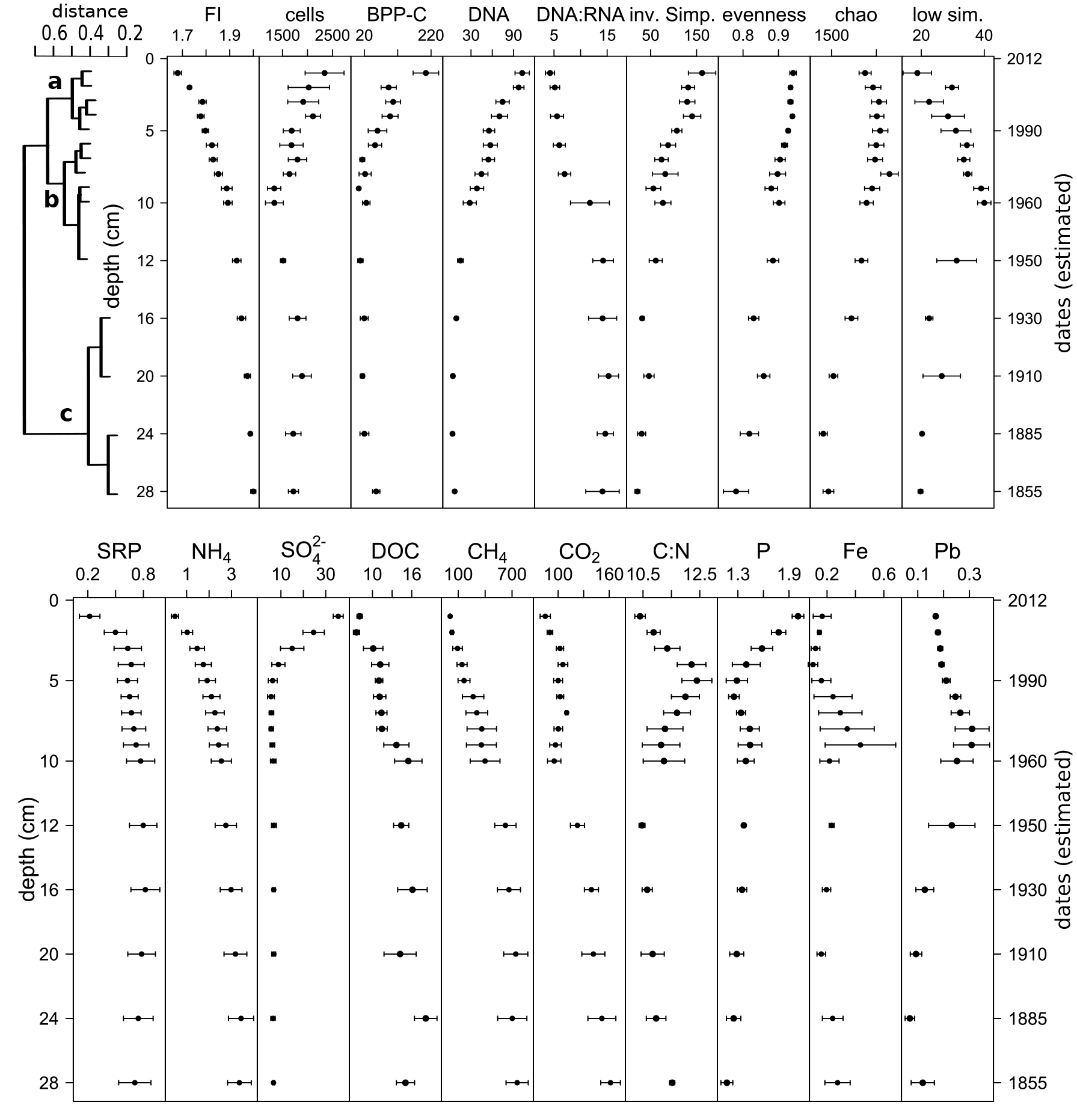
Depth profiles of a reduced set of sediment variables of Lake Stechlin at 30 *m* depth presented as the mean with standard error. The cluster dendrogram (average clustering) to the left presents the similarity of the total microbial community of each sampling depth and identifies the three depth horizons. A detailed version of all measured variables per individual core can be found in the supplemental material. Age values were adopted from ^137^Cs measurements. Units: cells [10^6^*ml*^−1^]; BPP-C/bacterial protein production in Carbon [*μg · ml^−1^d^−^1*]; DNA [*ng*]; low sim. [% of sequences]; SRP [*mg* · *l*^−1^]; *NH_4_* [*mg* · l^−1^]; *SO4^−^ [*mg ·* l^−1^*]; DOC [*mg · l^−1^*]; *CH_4_* [*μmol · l^−1^*]; *CO_2_* [*mmol · l^−1^*]; P [*mg · g*^−^1 dry weight]; Fe [mg · g^−^1 dry weight]; Pb [*mg · g*^−^1 dry weight]

Using a Chao estimate, the taxa diversity across the 60 samples was estimated to be 8545 (*SE* = 173) operational taxonomic units (OTUs). All a-diversity indices (inverse Simpson, evenness, and estimated Chao index) decreased with depth. The proportion of sequences that had no close known relative in the SILVA reference database (<93% sequence similarity by BLAST) was highest at 10cm depth (40 ± 4%).When we summarized the replicates and analyzed the community matrix globally using a cluster analysis, the sediment communities were grouped into three major clusters corresponding to the depth zones (0—5 cm, 5—14 *cm*, and 14—30 *cm)*, with a pronounced separation at 14 *cm* (Figure 1). Sediments at 15—30 *cm* depth were more similar in their community composition (> 65%) than in the upper two clusters (<50%). The community turnover (distance) follows a distance decay curve with increasing depth, approaching a distance of 1 (i.e., no shared taxa) for the lowest sediment layer compared to the surface layer (Table 1). When partitioning the diversity among sediment layers into richness and replacement effects, the effect of richness was proportional to increasing depth (R^2^ = 0.96, *F* = 230.8, *dF* = 11, see Figure S3), but it reached significance only at the deepest sampled layer (26—30 *cm* depth). Taxon turnover/replacement was consistently high and significant for multiple sediment depths within the first 14 cm (Table 1).

The OTUs that were the most influential in shifting the community structure across the 15 sediment depths were identified by calculating the corresponding species contribution to fi-diversity (Figure S4) [35]. The identities of these 96 “structuring” OTUs indicate an interrelationship of all three domains, with the most influential phyla being Euryarchaeota and Thaumarchaeota in the archaea and Chloroflexi, Proteobacteria, and Phycisphaerae in the bacteria. Redox-dependent groups (13% of the taxa could be clearly assigned to redox processes by their classification, e.g., Nitrospiraceae, Desulfobacteraceae, and Methylococcales) and redox irrelevant groups (eukaryotes and e.g., Bacteriovoraceae) structured the vertical microbial community. Out of these 96 “structuring” OTUs, we identified the OTUs that are significantly elevated in each of the three zones (Figure 2). While we see eukaryotic and bacterial lineages to be characteristic for the uppermost zone (cluster a in Figure 1), archaea and bacteria are elevated in the lowest zone (cluster c). For cluster b, only two structuring archaeal OTUs were identified. The residual OTUs from cluster b are significantly elevated either in the upper two clusters (mainly bacteria) or in the lower two clusters (mainly archaea). Out of the structuring OTUs, only one (Methylococcales) was significantly different in all three clusters (Figure 2).

**Figure 2.**
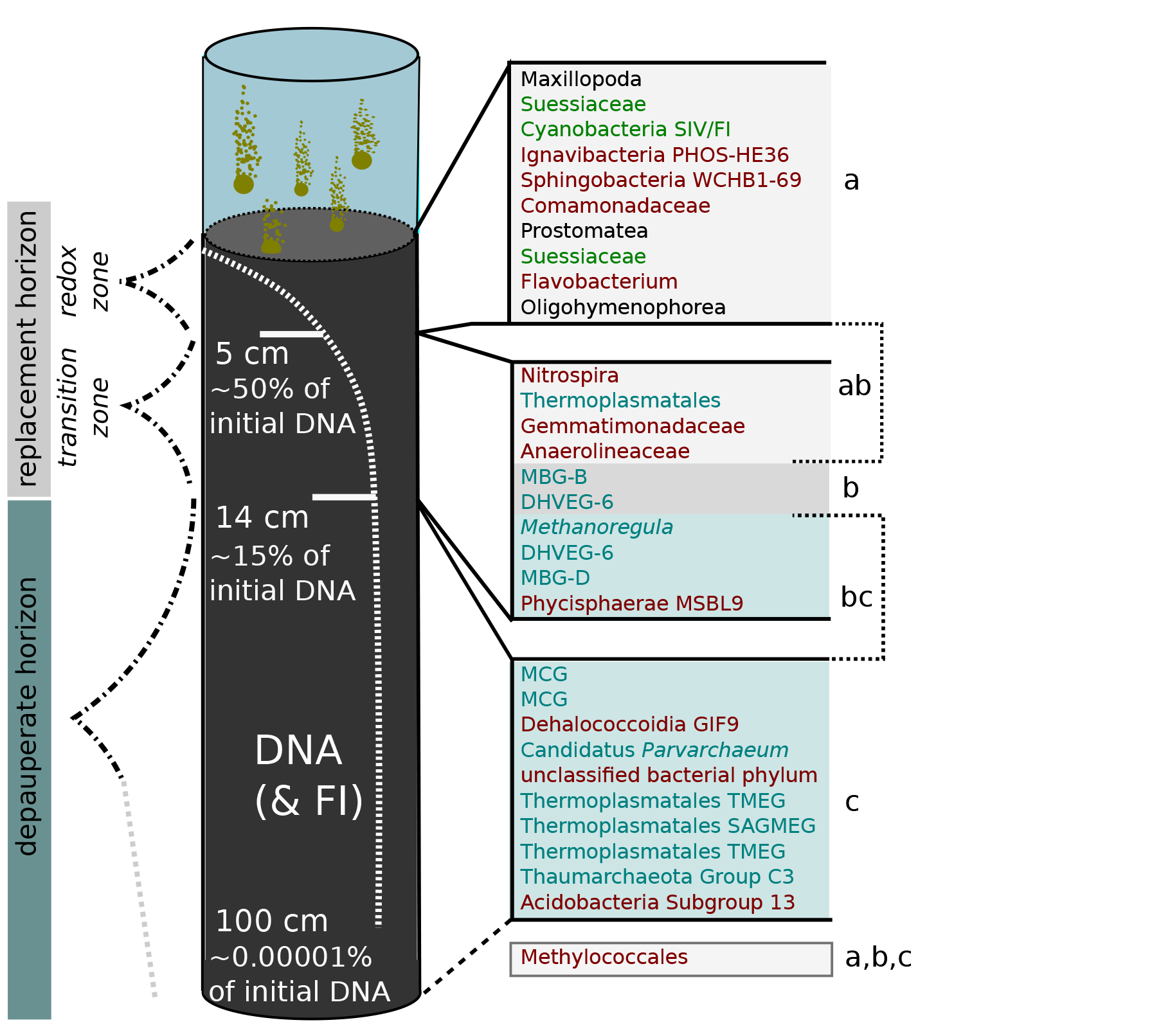
Overview on Lake Stechlin’s sediment structure. The cluster analysis separates three depth horizons: The *redox-stratified zone* (0—5 *cm*), which includes a thin layer of oxygen. A few fauna species exist in this zone, i.e., Nematoda, Gastrotricha, and microeukaryotes (e.g., Ciliophora), in addition to large numbers of highly active bacteria. Below 5 *cm*, where 50% of the DNA is already decomposed, the system enters the *transition zone*. This zone is situated below the sulfate-methane transition. Below 14 cm, we find the *depauperate horizon*, which extents in the deeper sediment, in which archaea dominate the community. In an extrapolation of the richness component of the community structure, the loss of richness would completely dominate (100%) the microbial community at 1 m depth (approx. 500 a). Following the decay curve of the DNA, 99.99999% of the DNA would be transformed at that depth. On the right side, the 10 most structuring OTUs (from Figure S4) are listed, which were significantly elevated in the corresponding horizon (only results with *p* < 0.01 in the TukeyHSD PostHoc Test, were included). The brackets ab and bc mark those OTUs that were elevated in the upper two or lower two zones, respectively. Only two OTUs were elevated in the transition zone. The grey box marks the single taxon that was significantly different in all three horizons. Taxon names are color coded according to their classification or phototrophy if applicable: phototrophic organism (green), eukaryotes (black), bacteria (red), archaea (blue).

**Table 1.**
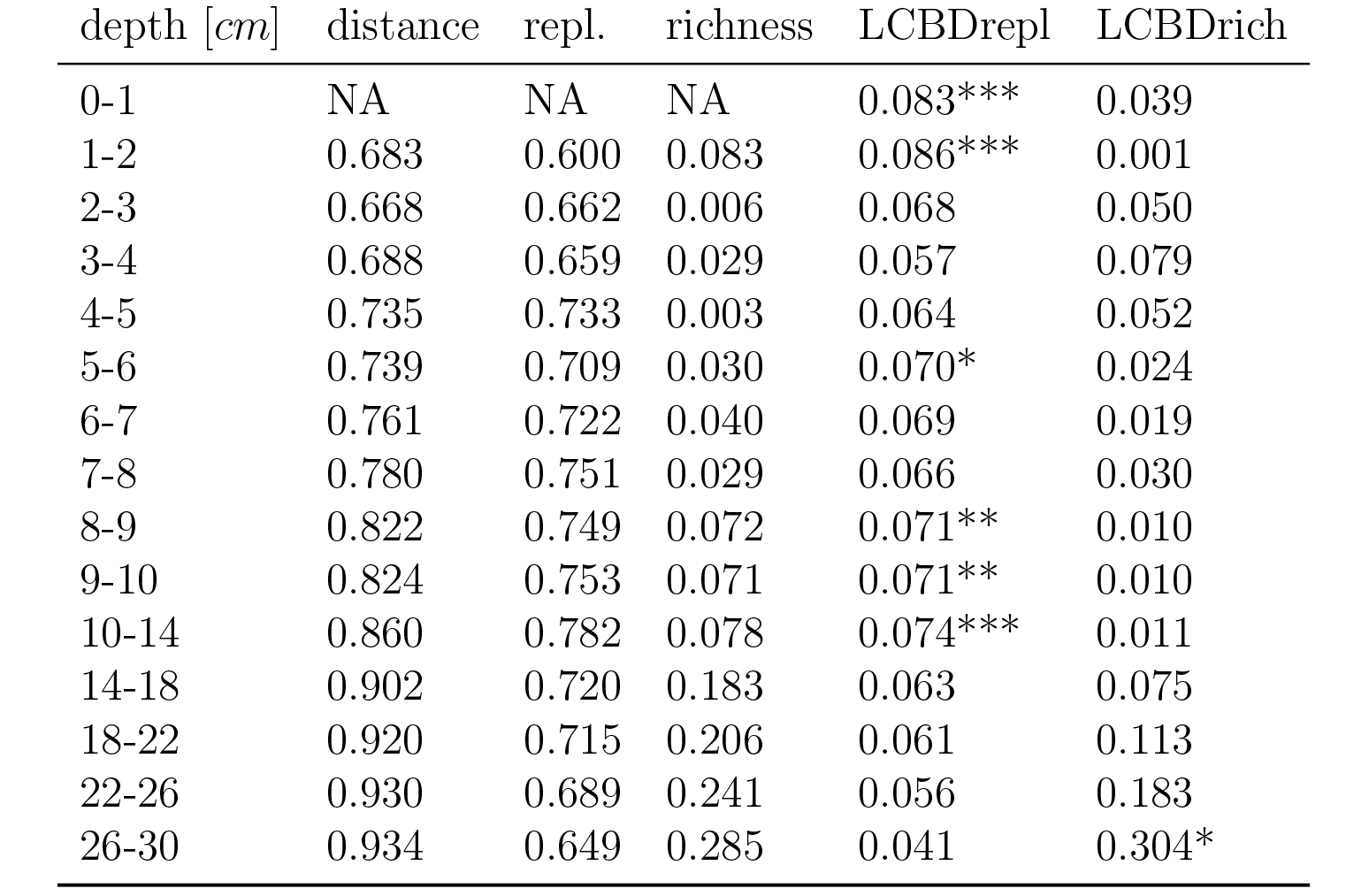
Jaccard based distance measures for all depths. Distance: total Jaccard based distance to the uppermost depth [0-1 cm]; repl.: replacement component of the Jaccard distance; richness: richness component of the Jaccard distance; LCBDrepl: local contribution of the replacement component; LCBDrich: local contribution of the richness component

| depth [cm] | distance | repl. | richness | LCBDrepl | LCBDrich |
| --- | --- | --- | --- | --- | --- |
| 0-1 | NA | NA | NA | 0.083*** | 0.039 |
| 1-2 | 0.683 | 0.600 | 0.083 | 0.086*** | 0.001 |
| 2-3 | 0.668 | 0.662 | 0.006 | 0.068 | 0.050 |
| 3-4 | 0.688 | 0.659 | 0.029 | 0.057 | 0.079 |
| 4-5 | 0.735 | 0.733 | 0.003 | 0.064 | 0.052 |
| 5-6 | 0.739 | 0.709 | 0.030 | 0.070* | 0.024 |
| 6-7 | 0.761 | 0.722 | 0.040 | 0.069 | 0.019 |
| 7-8 | 0.780 | 0.751 | 0.029 | 0.066 | 0.030 |
| 8-9 | 0.822 | 0.749 | 0.072 | 0.071** | 0.010 |
| 9-10 | 0.824 | 0.753 | 0.071 | 0.071** | 0.010 |
| 10-14 | 0.860 | 0.782 | 0.078 | 0.074*** | 0.011 |
| 14-18 | 0.902 | 0.720 | 0.183 | 0.063 | 0.075 |
| 18-22 | 0.920 | 0.715 | 0.206 | 0.061 | 0.113 |
| 22-26 | 0.930 | 0.689 | 0.241 | 0.056 | 0.183 |
| 26-30 | 0.934 | 0.649 | 0.285 | 0.041 | 0.304* |

For a general overview on the median appearance of eukaryotic, bacterial, and archaeal taxa at single depth layers, see Figure S1. Globally, archaea, bacteria, and eukaryotes all exhibited pronounced vertical changes. Their sequence proportions (A:B:E) shifted from 10:70:20 at 0 *cm* to 50:50:0 at 10 *cm* and 60:40:0 at 30 *cm* depth (Figure 3A). The eukaryote pattern was correlated with the total DNA concentrations in the sediment (r = 0.869), and it decayed exponentially with depth. We were able to predict DNA concentration using a multiple linear regression as a function of the occurrence of eukaryotes (75.6% of the variation) together with bacteria (10.0% of the variation; model: R^2^ = 0.856,*p* < 0.001, Figure S5). A multivariate ordination of all the samples confirmed a strong vertical gradient in the community structure, which was reflected in the distance between the surface and deep sediments on axis 1 (Figure 3B, Mantel correlation: *r* = 0.735,*p* < 0.001). We were able to significantly recover the three depth zones (Figure 3B, PERMANOVA: *F* = 12.3,*p* = 0.0005), and there was an obvious reduced variance in cluster c compared to the other clusters (betadispersal analysis: Tukey’s Honest Significant Differences between groups, *p* <0. 01), confirming the higher similarities seen in the previous cluster analysis. The overall community structure was correlated with both present (Mantel correlation: *r* = 0.527,*p* < 0.001) and past (*r* = 0.459,*p* < 0.001) parameters, which were nearly orthogonal in ordination. To elucidate this further, we investigated whether the past parameters may match with the richness component of the community composition, and correspondingly, whether the present parameters are correlated to the replacement component of the microbial community. Thus, we partitioned the whole dataset into replacement and richness matrices and correlated these to the present and past parameters by employing a fuzzy set analysis. The richness community submatrix was strongly correlated with the past parameters (two-dimensional fuzzy set ordination with the first 2 axis of the PCA, *r* = 0.99). The replacement community submatrix was correlated with the present parameters (one-dimensional fuzzy set ordination with the first axis of the PCA, *r* = 0.65,*p* < 0.001).

**Figure 3.**
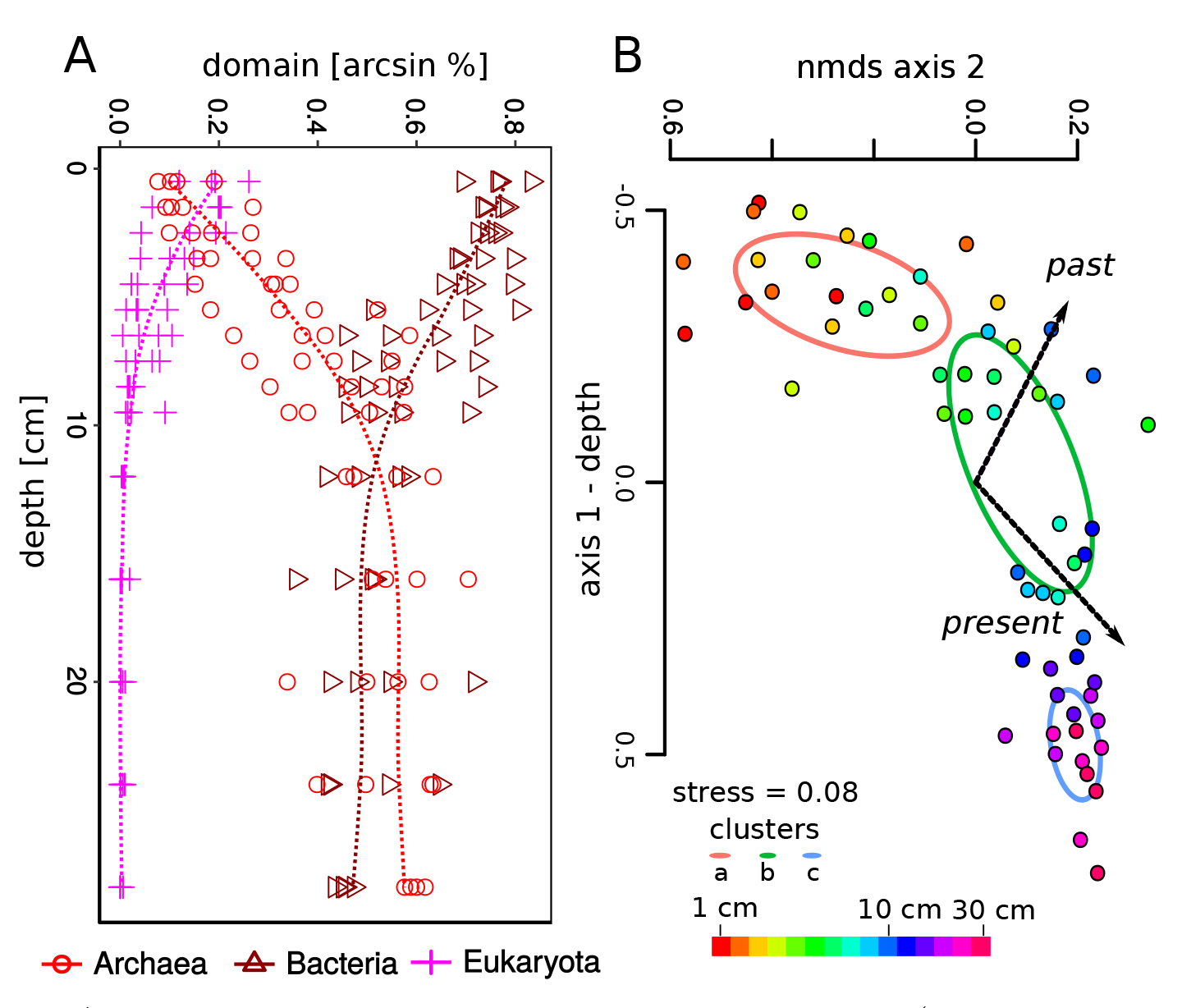
A. Depth profiles of the microbial community (eukarya, bacteria, and archaea) presented as relative proportions to each other, which was determined by relative pyrosequencing reads per microbial fraction. B. NMDS ordination of the vertical sediment microbial community structure. The clusters from Figure 1 are presented as standard deviation around the group centroid. The color scale of the dots represents sediment depth.

## Discussion

Comprehensive studies of the microbial communities and physic-chemical characteristics of freshwater sediments are scarce, and general concepts are often transferred from marine systems without validation. Often salinity and significant higher sulfate concentrations are responsible for major differences between both systems [36,37]. We address fundamental questions regarding the structure and organization of freshwater sediments and establish a structural model that can potentially be applied to other aquatic sediments.

### Organization of freshwater sediments

Although constant microbial colonization of surface sediments occurs via sinking OM, the sediment habitat is considered to be an autonomous system in terms of species diversity and community structure [17, 18], with the exception of surface supply of planktonic OM, including decaying eukaryotes (Figure S4). Indications that a high species turnover, as suggested by our results, may be a common feature of vertical sediment profiles have been reported for bacterial taxa in coastal marine sediments [38], for marine archaea and bacteria [39], and for freshwater archaea [20]. Previous freshwater sediment studies that found a more moderate species turnover were restricted by the use of low-resolution methods [15, 20, 40]. Moreover, most studies have focused on either bacteria or archaea and not on all three domains simultaneously, and none of the previous marine or freshwater studies partitioned-diversity into richness and replacement components, which allowed us to look at both buried and active taxonomic signals.

Our results pointed to three depth clusters. Based on the high replacment (Table 1) and measurable microbial production in the first two depth clusters (cluster a, b) (Figure 1), we decided to classify them into one overarching horizon, which we called the replacement horizon. Correspondingly, we classified the lowest depth cluster (cluster c) into the depauperate horizon, based on the loss in richness measures and constancy in most sediment parameters (Box 2).

##### Box 2. *Sediment zonation according to taxonomic clustering, fi-partitioning, and context data*

- I.The replacement horizon (1—14 *cm*|*ca*. 0—60 *a*) is subdivided into two zones, delineated by the sulfate-methane transition, which falls approximately within or correlates with a local prokaryotic cell maximum.

- Cluster a) redox-stratified zone (0—5 *cm*|*ca*. 0—20 *a*). This zone encompasses the redox-stratified zone and the typical sequence of electron acceptors from oxygen to sulfate. 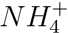 and SRP increases. It is characterized by high microbial activity, cell numbers, taxa turnover, spatial variability, low DNA:RNA ratios, and low FI values (1.7—1.8). Potentially 50% of the sedimented organic matter is metabolized in this horizon, aligning with the rapid decrease of eukaryotes. Bacteria dominate this horizon.
- Cluster b) transition zone (5—14 *cm*|*ca*. 20—60 *a*). This zone is characterized by strong gradients. Microbial cell numbers drop off and activity decreases, diversity decreases, the taxa turnover stays high, the DNA:RNA ratio doubles, and FI values increase (1.8—1.9). Methane concentrations rise, and *CO_2_* has a local minimum. Potentially, another 35% of the sediment organic matter is turned over in this horizon. Eukaryotes approach zero, bacteria decline, and archaea rise rapidly in their community contribution. Here, we also see a maximum of taxa with no close relatives to known database entries.

- II.The depauperate horizon (14—x *cm*|*ca*. > 60 *a*) is very distinct from horizon I, and it is unclear how far this horizon reaches. It appears to be archaea-dominated and is characterized by a loss in richness and a shift toward the dominance of single taxa.

- Cluster c) (14—30 *cm*|*ca*. 60—150 *a*). The zone is characterized by constancy in most parameters, while we see significant taxon richness effects and marked drops in spatial variability, evenness, and diversity. Cell numbers and activity stay consistently low, while DNA:RNA ratios and FI values (2.0) stay consistently high. *CH_4_* and *CO_2_* approach saturation. Archaea are dominant.

**The replacement horizon** Bacterial activity was highest in the redox-stratified zone (cluster a), which is where most of the settled OM is available. As sulfate is depleted at 5 *cm*depth, most redox processes will take place above this. The majority of freshwater sediment studies examine this zone in great detail (e.g., [4,5, 10]), including the identification of redox processes at the millimeter scale [41,42]. Highly active decomposition processes lead to high prokaryotic cell numbers close to the sediment surface (e.g., [5,24,43], as well as a constantly lower abundance in deeper horizons [15]. There was also an initial loss of many taxa in the highly active oxycline-indicated in our data by an outlier to the richness community component (Figure S3)-which was potentially intensified by active grazing through, for example, ciliates and copepods. The sulfate-methane transition marks the border of the transition zone (cluster b). This border can harbor important methane oxidation processes [44]. At 10 *cm*, the *CO_2_* minima might indicate the beginning of hydrogenotrophic methanogenesis (Figure 1). [45] described this pathway for profundal Lake Stechlin sediments. In this second subzone, the archaeal contribution rises constantly, while bacteria and eukaryotes decrease and the general activity measures decline rapidly. As indicated by the DNA measurements, more than 85% of the settled organisms were decomposed within this replacement horizon, which spanned approximately 60 years. The decay of DNA, in combination with an enrichment of fulvic acids with increasing depth (FI values), confirms the assumption that DNA can serve as a proxy for the buried OM. Thus, there is not only a gradient of electron acceptors in the replacement horizon but also a gradient of OM quality. This might again facilitate the stratification of active microbial taxa with depth since OM can simultaneously serve as an electron donor and acceptor [46,47] and because OM quality is known to modulate microbial redox processes [48], with apparent consequences for carbon turnover rates [33,47].

**The depauperate horizon** This horizon (cluster c, > 14 cm) is fairly distinct in many parameters; it features high methane and CO2 concentrations, a dominance of archaea, and richness impoverishment. In this zone, the theoretical model of a hierarchical, nested system is clarified (9% vs. 2% relative nestedness in the replacement horizon). At 14 *cm*, the DNA:RNA ratio doubled and archaea replaced bacteria as the dominant microorganisms. We believe this reflects an increase in the number of microbes entering a stationary state below this depth, where cell maintenance overrules cell synthesis due to the low availability of terminal electron acceptors. This is analogous to what has been suggested for cells in low-energy marine environments [26,28,49].

The variability in community composition is very low across replicates in this horizon. The system is characterized by the gradual disappearance of taxa with burying age and a richness component steadily increasing to more than 20% (Table 1Table 1). If the richness component further increased linearly, it would be the sole factor in structuring the community composition in 1 *m* sediment depth (the total sediment depth of Lake Stechlin is 6 *m*). It is intuitive that the richness component may be a function of the burying time and that it represents the fading signal of preserved organisms. However, it is not clear why it does not follow an exponential decay function that is analogous to that for DNA.

**Potential causes of the high taxa replacement** In our study, many parameters changed rapidly, especially in the replacement horizon (DNA, FI, BPP, electron acceptors), propagating microbial taxa turnover. This could be potentially caused by several different mechanisms: **a)** Cellular turnover: taxa replacement is potentially caused by cell synthesis, lysis, and recycling of dormant cells, which are assumed to be high in sediments [50,51]. In particular, viral lysis of cells is supposed to be very important in sediments [52,53]. We found indications for cellular recycling caused by the predatory Bacteriovoracaceae (cf. [54]), which is one of the structuring bacterial lineages identified in Figure S4. Another potential mechanism-one that may be most important in the depauperate horizon is differential cell replication. The resources for cell maintenance and growth should depend on cell size and complexity, which means that small cells, such as nanoarchaea (e.g., Candidatus Parvarchaeum), should have a selective advantage as they can continue to grow under conditions that cause other cells to switch to cell maintenance. This could be one explanation for the observed drop in evenness in the depauperate horizon. **b)** Random appearances: Replacement might also be caused by the random appearance of taxa due to the disappearance of others. Since high-throughput sequencing (HTS) produces relative data, it may superimpose proportions over quantities. For example, the initial decay of eukaryotes may have opened a niche for previously hidden rare taxa. Further, if there was no growth in the sediment, lineages that are potentially better suited for long-term survival than others would appear, such as spore-forming bacteria (Firmicutes). However, we (and others: [26,31]) did not observe an enrichment of this lineage in the HTS data with depth (data not shown). In addition, the use of replicate cores in combination with our conservative stripping (see the Method section) should have removed most of the random effects. The more or less constant cell numbers with small local maxima, and the observed shifts in the evenness support a rather non-random stratification of sediment communities including cell replication. While the cellular reproduction probably approaches stagnation for most microbes in the depauperate horizon, the slowly shifting redox conditions during different seasons and across years may be conducive for the colonization of the replacement horizon by different niche specialists. The low sedimentation rate of Lake Stechlin (ca. 2 mm per year, as determined by ^137^Cs-dating at the Federal Office for Radiation Protection, Berlin [courtesy of U.-K. Schkade], or 0.4—2.1 *g* · *m*^−2^·*d*^−1^, as determined by sediment traps [55]) may mitigate this stratification, and the scale of the horizons may look different in systems with higher or lower sedimentation rates.

### Burial processes and microbial activities

The microbial sediment community appears to be highly indigenous, yet burial processes take place simultaneously, and the microbial community itself may eventually be buried. The buried DNA and organisms seem to preserve historical plankton communities, which should resemble the past conditions of the lake ecosystem [31,56, 57]. These past environmental conditions are partly preserved as particulate matter, which is relatively conservative. We were able to show that these conservative past parameters still correlate with the overall community. Although past studies have shown that sediment parameters influence community patterns in marine systems [39], the respective context data were not separated into present and past, nor was the microbial community separated into richness and replacement components. In this context, it is interesting that the richness component can be well explained by the first two principal components of the past parameters. Since the replacement component could not be fully mapped by the present parameters in our study, we can assume that sources of variation other than environmental parameters, in particular, biotic taxa interactions, are relevant. Strong biotic interactions have been identified in a vertical profile of a meromictic lake with a comparable chain of redox processes as they occur in sediments [58]. Deep sediment layers may offer specific energy-poor microniches favoring a high variety of syntrophic microorganisms [59,60]. The Dehalococcoidales (Chlorflexi [61,62]) and the Miscellaneous Crenarchaeotic Group (MCG, [63,64]) are promising candidates for such hybrid forms of energy harvesting and are both among the most influential lineages in our data (Figure S4). In our data, the MCG co-occur with Dehalococcoidales, similar to their presence in Lake Baikal’s (> 1500 m water depth) methane hydrate-bearing sediment [65]. Another indication that biotic interactions are important in Lake Stechlin sediments is the appearance of the Candidatus Parvarchaeum as a structuring lineage (Figure 2). Thislineage can exhibit a cell-to-cell coupling, allowing for thermodynamic processes that 297 would otherwise not be possible [66].

### Patterns of freshwater archaea

The fact that archaea can be very numerous in freshwater sediments and even dominate microbial communities is a rather new discovery, and hard facts based on comparative studies are still lacking. Their recovery rate in relation to bacterial sequences or cell numbers varies between 3-12% ( [67], cells), 5-18% ( [20], qPCR), and 14-96% ( [14], qPCR), depending on the lake, sampled sediment horizons, and methods employed. In most cases, only surface sediment samples have been considered (e.g., [68], 1% of cells), and the few studies involving vertical profiling to date are ambiguous in finding an archaeal depth gradient. Our results and the cell counts of [67] in Lake Biwa (Japan) point to an increase in the proportion of archaea with sediment depth. However, the results obtained by quantitative PCR for Lake Taihu (China; [14]) and Lake Pavin (France; [20]) fail to show such a relationship. Archaea have, on average, compact genomes [69] and a lower ribosomal copy number than bacteria [70], which may lead to underestimates of the actual archaeal abundance. Similar to our results, [20] found three sequential depth clusters in the archaeal community structure within the first 40 *cm*, defining an intermediate layer between 4 and 12 *cm*.Next to well described methanogenic archaea we, similar to the findings of [20], mainly recovered archaeal lineages with no clear functional assignment thus far, i.e., primarily the MCG (potentially methanogenic, [64]) and the Marine Benthic Group D (MBG-D). Both groups are among the most numerous archaea in the marine sub-seafloor, and they are suspected to metabolize detrital proteins ( [19], discussed in more detail below). Interestingly, we also identified a MCG-B as structuring OTU for the transition zone (Figure 2), a group which was recently described as eukaryotic progenitor from a hydrothermal vent field (Lokiarchaeota, [71]). Also, several MCG OTUs belonged to the top structuring taxa. MCG was recently named as Bathyarchaeota by [72] for its deep-branching phylogeny and its occurrence in deep subsurface environments -environmental conditions that our cores (30 m water depth and 30 *cm*length) did not meet. Our results suggest that the specific niche adaptation of these microbes is not necessarily related or restricted to the deep biosphere but rather to a cellular state of “low activity” [73]. In this context, it is interesting that single MCG OTU sometimes dominated the community in the deep horizons (up to 34% in core D at 26—30 *cm*), resulting in a reduced overall evenness and a shift of the residual taxa to the rare biosphere, contrasting the potential random effects as discussed above. Another intriguing observation is the considerable overlap of archaeal and partially bacterial lineages between our study and deep-sea environments. Consequently, typical marine lineages (e.g., archaea in Figure S1: MGI, MCG [Bathyarchaeota], MHVG, DHVEG-1, -6 [Woesearchaeota], DSEG, MBG-A, -B [Lokiarchaeota], -D, -E) are not as “marine” or as “deep-sea” as previously thought. Given the high cost of deep-sea research [28], freshwater sediments might literally pose a row-boat alternative for research questions targeting these “remote” and “extremophile” microorganisms.

## Summary and Conclusions

Our results point toward a steady-state chemostat-like environment with a highly stratified indigenous microbial community in freshwater sediments. Sediments are neither a one-way system for the burial of organic matter nor purely composed of redox-active microorganisms. Both processes take place alongside a vertical gradient of electron acceptors and decomposing OM of decreasing quality. The microbial community is highly structured into distinct zones, matching most sediment parameters. Both the microbial community and the sediment parameters can be divided into components relevant for burial and past conditions as well as for recent carbon turnover processes and their context data. Biotic interactions are likely to play an important role, and we were able to identify important sediment taxa for each horizon. We put a spotlight on the largely unexplored freshwater sediments and confirmed earlier findings that were previously described only for marine sediments, such as the importance of marine archaeal lineages and the introduction of a depauperation zone in which the burial process becomes increasingly important.

## Materials and Methods

### Sampling site and sampling procedures

Lake Stechlin (latitude 53°10 N, longitude 13°02 E) is a dimictic meso-oligotrophic lake (maximum depth 69.5 *m*; area 4.23 *km^2^*) in northern Germany that has been the subject of more than 55 years of research [74]. Sediment cores were extracted from the southern bay of the lake. Four adjacent sites were sampled to account for spatial heterogeneity in the sediment (Sites A-D, Fig. S6). Pore water was collected by four in-situ dialysis samplers, so-called “peepers” [75], which were deployed for 14 days using a frame (1 *m*^2^). Shortly before retrieving the peepers, four sediment cores were taken from each site with Perspex tubes (inner diameter: 6 or 9 *cm*; length: 60 *cm*) using gravity corers (UWITEC™, Mondsee, Austria) at 30 *m* water depth (aphotic depth) on two subsequent days (March 26 and 28, 2012; peepers were retrieved on April 1, 2012). Two sediment cores (6 *cm*diameter) were stored in the dark at 4°C until oxygen penetration depth was measured within the next 4 hours (see below). The sediments of the 9 *cm*cores were sliced directly into 1 *cm*layers for the uppermost 10 cm, then in 4 *cm*layers for sediment depths of 10—30 *cm*. One core was used for the analysis of the total sediment and the other was used for pore water, gas, and microbial analyses (see below). In May 2014, 24 additional cores were taken to determine the age-depth correlation using the cesium 137 technique.

### Maximum oxygen penetration depth

Two initial cores were carefully transferred into 20 *cm* short cores without disturbing the sediment surface. The short cores were kept cool (4°C) until measurements were taken. Oxygen microprofiles were performed using two Clark-type microelectrodes (OX50 oxygen microsensors, Unisense, Aarhus, Denmark) with a 50 μ*m* glass tip. SensorTracePro 2.0 software (Unisense) was used for data storage. The electrodes were calibrated by two-point calibration. For each core, we measured at least four profiles. The sediment-water interface was defined as the point where the oxygen depletion shifted from linear to non-linear [76].

### Pore water analysis

The sampled sediment horizons were centrifuged (13,250 g for 10 min) to retrieve pore water (filtered through rinsed 0.45 μ*m* cellulose acetate membranes, Roth, Germany) for immediate analysis of the dissolved organic carbon (DOC) and FI. DOC was measured as non-purgeable organic carbon with an organic carbon analyzer (multi N/C 3100, Analytic Jena AG, Jena, Germany). FIwas measured following the protocol of [34]. Peeper samples were analyzed for concentrations of SRP and ammonium (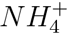), dissolved iron (Fe^2+/3+^), manganese (*Mn*^2^+), chloride *(*Cl*^−^)*, nitrate (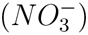), and sulfate (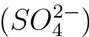), following DIN EN ISO 10304-1. SRP and NH+ were photometrically determined using a segmented flow analysis (SFA, Skalar Sanplus, Skalar Analytical B.V., De Breda, Netherlands). Dissolved iron and manganese levels were determined by AAS (Perkin Elmer 3300, Rodgau-Juegesheim, Germany), and analyses of the dissolved anions nitrate and sulfate were conducted by ion chromatography (IC, Shimadzu Corporation, Japan).

### Total sediment analysis

Sediment water content was analyzed by drying at 85°C until mass was constant. A subsample was used to determine the organic matter content (4 h at 550°C) of the sediment. The metal concentrations were determined by ICP-OES (iCAP 6000, Thermo Fisher Scientific, Dreieich, Germany) after aqua regia digestion in a microwave oven (Gigatherm, Grub, Switzerland), and total phosphorus (TP) was determined by CARY 1E (Varian Deutschland GmbH, Darmstadt, Germany) after *H_2_SO_4_/H_2_O_2_* digestion (150°C, 16 h). CNHS content was determined using aliquots of dried matter in a vario EL system (Elementar Analysensysteme GmbH, Hanau, Germany).

### Gas chromatography

From each depth, 2 ml of sediment was transferred into 10 ml vials filled with 4 ml of distilled water. Samples were fixed with mercury chloride (final conc. 200 *mgl*^−1^), sealed, and stored in the dark at 4°C until analysis. Concentrations of *CO_2_, CH_4_, and N_2_O* were measured by gas chromatography (Shimadzu GC-14B, Kyoto, Japan) using the headspace technique described in [77].

### Bacterial protein production

Bacterial biomass production was determined via ^14^C leucine incorporation at in situ temperature under anoxic conditions [78] using a modified protocol [79]. 500 *μl* of sediment was diluted 1:1 with sterile filtered supernatant water and incubated with 14C-leucine (Hartmann Analytics, Braunschweig, Germany; specific activity:306 *mCi mmol*^−1^, diluted with cold L-leucine to a final concentration of 50 *μmoll^−1^*).Incubations were stopped after 1 h, extracted, and measured in a liquid scintillation analyzer (TriCarb 2810 TR, PerkinElmer Inc., Germany). Disintegrations per minute were converted to pmol leucine *ml^−1^ day*^−1^, assuming a two-fold intracellular isotope dilution [80,81].

### Cell counting

Sediment subsamples for cell counting were immediately fixed with ethanol (50% v/v final concentration). Prior to analysis, samples were shaken for 1 h at 700 rpm on a thermoshaker and were sonicated three times for 20s at 5—*6W* (Branson Sonifier 150, Danbury, USA). Cells were stained with a SYBR Gold staining solution diluted to 1:1,000 (Molecular Probes, Eugene, USA) and were counted with an epifluorescence microscope (Zeiss, Axio Imager. Z1, Jena, Germany).

### Nucleic acid extraction and sequencing

To determine the DNA:RNA ratio, we extracted total nucleic acids using a phenol-chloroform protocol from 200—400 *μl* sediment, as described by [82]. The DNA:RNA ratio was measured via fluorometry using selectively binding dyes (Broad range dsDNA and broad range RNA assay Kit, Life technologies, Darmstadt, Germany) developed for the Qubit 2.0 (Life technologies, Darmstadt, Germany). A second extraction served as template for the sequencing and determination of the total DNA content. A defined sediment subsample (350*μl*) from each depth was lyophilized prior to DNA extraction. We used the “Alternative Protocol for Maximum DNA yields” of the UltraClean^®^ Soil DNA Isolation Kit (MoBio Laboratories Inc., Carlsbad, USA). The quality of the DNA and the presence of putative environmental (small) DNA in 12 representative samples from 12 depths were verified with a microgel electrophoresis system (DNA High Sensitivity Kit, Bioanalyzer, Agilent, USA, see Fig. S7). A total of 5—20 *ng* of DNA, as measured by NanoPhotometer P300 (Implen, Schatzbogen, Germany), served as the template for PCR amplification (Herculase II system, Life Technologies) using a single universal primer system (926F, 1392R, [83]). The primer pair employed is one important feature of our study in that it detects all three microbial domains (archaea, bacteria, and eukaryotes) in freshwater systems [58,84]. PCR products were purified with AMPure Beads (Beckmann Coulter, Brea, USA) and quantified and pooled using a PicoGreen assay (Life Technologies, Carlsbad, USA). High-throughput sequencing was performed in a Roche 454 GS Junior benchtop sequencer (Hoffmann-La Roche, Basel, Switzerland) at the Berlin Center for Genomics in Biodiversity Research. Raw sequence data were deposited in ENA (accession PRJEB14189).

### Data processing

454 sequencing data were processed using Mothur (version 1.33.0) following the guidelines of the Mothur SOP (http://www.mothur.org/wiki/454_SOP, accessed 02/2014) with the following modifications: (i) for quality trimming, we used a sliding window approach with a relaxed threshold (window size: 50, quality cut-off: 27), and (ii) the alignment step used SINA (version 1.2.11; [85]) against the SILVA v.111 non-redundant SSU reference database. A total of 396,000 reads (49% of raw sequences) were retained. The sequences were clustered into OTUs at 97% sequence similarity. A representative sequence from each OTU was used for taxonomic classification with the least common ancestor method in SINA, using 0.7 as a setting for minimum similarity as well as for lca-quorum. The classified OTU abundance matrix served as the basis for all subsequent statistical analyses. The percentage of sequences with low similarity (<93%) to the next reference sequence was determined by submitting the fasta files to SILVA NGS [86].

### Statistics

All measured environmental parameters were compiled in a matrix and imported into R (http://cran.r-project.org/, version 3.2.2). We replaced two outliers (FI: replicate 4 cm, total phosphorous: replicate 14 cm) with the mean values of the three other sediment cores. Similarly, the 30 *cm*peeper data from replicate core B were missing and replaced by the mean of the residual replicates. For statistical analysis, the relative proportions of archaea, bacteria, and eukaryotes were arc-sin transformed. For the multiple regression analysis on the declining DNA concentrations, we removed one value (replicate 3, 22 *cm*) to meet the normal distribution criteria of the residuals. Sufficient normal distribution was confirmed by a QQ plot and Shapiro-Wilks test, *p* = 0.183; Cook’s distance was not violated in any case. We categorized the environmental parameters into present (*CH*_4_, *CO*_2_, DOC, BPP, SRP, 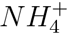, 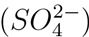, *Cl*^−^, *Fe*^2^+^/3^+, Mn^2^+, FI, and RNA:DNA) and past (TC, TN, dry-weight, TP, TS, TH, Al, As, Ca, Cu, Fe, Mg, Mn, Pb, Ti, and Zn) (see Box 1) and assessed the sample variation for each subset by a centered, scaled principal component analysis (PCA, see Fig. S8). The resulting most explanatory PCA axes, which explained 54% (present parameters) and 41% (past parameters) of the sample variation, were used in the community statistics (see below). DNA and cell numbers were not categorized due to their ambiguous nature; Al was used as a substitute for Mg and Ti due to their high degree of correlation (*r* > 0.95); N_2_O, 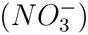, Cd, and Co were excluded due to their very low values, i.e., near or below the detection limit in all the samples.

### Community statistics

Random effects were initially excluded from the OTU matrix by removing all OTUs present in only one sample, regardless of the number of reads. The random effects are expected to be very high in sediments due to the burial of random organic matter (e.g., caused by the droppings of a bird, tourist activities, rainfall) and thus a large amount of rare taxa. This reduced the number of OTUs from 29,228 to 9,581 but did not influence the sample distances (Mantel test with Hellinger distances: r = 0.993,p < 0.001). Diversity indices were calculated using the vegan package [87] for R, with a community matrix that was rarefied to the lowest number of reads (642) present in a sample. The rarefied matrix was highly correlated to the initial matrix (Mantel test with Hellinger distances: *r* = 0.923,p < 0.001). Nonmetric multidimensional scaling (NMDS) and Mantel tests were based on a Hellinger-transformed OTU matrix with Euclidean distances. The PCA scores from the past and present parameters (see Box 1) were fitted into the NMDS, and their correlation with the underlying distance matrix was tested with a Mantel test. We used a fuzzy set ordination [88] to test for the influence of the past and present parameters on the separated richness and replacement community components. For this, we partitioned fl-diversity into richness and replacement components using indices from the Jaccard family, following [89] and the functions provided by [35]. In order to identify general vertical patterns, we used a sum table to increase the resolution and thus avoid an artificial increase in turnover versus the richness/nestedness structure due to sampling effects. The sum table was generated by summing up 2,000 sequences per depth, if applicable. The final sum table was rarefied to the lowest number of sequences in the depth profile (5,987 reads). The cluster analysis (UPGMA clustering based on Kulczynski distance, Fig. 1) was also done on the sum table. The depth-dependent nestedness [29], richness component, replacement component, SCBD, and LCBD were calculated as described in [35]. We note that the nestedness index [29] is dependent on the sample size, and so we refer to it as “relative nestedness.”

## Acknowledgments

We thank Michael Sachtleben and Federica Pinto for their help during sampling. We also thank Elke Mach, Uta Mallok, Christiane Herzog, Hans-Jiirgen Exner, Grit Siegert, Susan Mbedi, Kirsten Richter, Camila Mazzoni and Andreas Kleeberg for their assistance and measurements in the laboratory and Henrik R. Nilsson for comments on this manuscript. This is publication 40 of the Berlin Center for Genomics in Biodiversity Research. This work was funded by the Leibniz Association Pakt for Research and Innovation project “Climate-driven changes in biodiversity of microbiota-TemBi” (SAW-2011-IGB-2) and DFG GR1540/15-1 and 23-1. We thank U.-W. Schkade (Federal Office for Radiation Protection, Berlin, Germany) for the dating of the Lake Stechlin sediments.

